# Citizen Science for Mining the Biomedical Literature

**DOI:** 10.1101/038083

**Authors:** Ginger Tsueng, Steven M. Nanis, Jennifer Fouquier, Benjamin M Good, Andrew I Su

## Abstract

Biomedical literature represents one of the largest and fastest growing collections of unstructured biomedical knowledge. Finding critical information buried in the literature can be challenging. In order to extract information from free-flowing text, researchers need to: 1. identify the entities in the text (named entity recognition), 2. apply a standardized vocabulary to these entities (normalization), and 3. identify how entities in the text are related to one another (relationship extraction). Researchers have primarily approached these information extraction tasks through manual expert curation, and computational methods. We have previously demonstrated that named entity recognition (NER) tasks can be crowdsourced to a group of nonexperts via the paid microtask platform, Amazon Mechanical Turk (AMT); and can dramatically reduce the cost and increase the throughput of biocuration efforts. However, given the size of the biomedical literature even information extraction via paid microtask platforms is not scalable. With our web-based application Mark2Cure (http://mark2cure.org), we demonstrate that NER tasks can also be performed by volunteer citizen scientists with high accuracy. We apply metrics from the Zooniverse Matrices of Citizen Science Success and provide the results here to serve as a basis of comparison for other citizen science projects. Further, we discuss design considerations, issues, and the application of analytics for successfully moving a crowdsourcing workflow from a paid microtask platform to a citizen science platform. To our knowledge, this study is the first application of citizen science to a natural language processing task.

## II. Background

Biomedical research is progressing at a rapid rate (Bornmann & Mutz, 2015). The primary mechanism for disseminating knowledge is publication in peer-reviewed journal articles. Currently there are over 25 million citations indexed in PubMed (“PubMed Help”, 2016), the primary bibliographic index for the life and health sciences developed and maintained by the US National Center for Biotechnology Information. PubMed is growing by over one million new articles every year.

Despite the exponential growth of the biomedical literature (Lu, 2011), accessing the accumulated knowledge that it contains remains a difficult problem. Journal publications are primarily in the form of free text, a format that is difficult to query and access. This problem is especially pronounced for biomedical research articles because of the imprecise way that language is used to refer to important biomedical concepts. For example, the acronym PSA has been used to refer to a number of different human genes, including “protein S (alpha)” (Ploos van Amstel et al., 1990), “aminopeptidase puromycin sensitive” (Osada, Sakaki and Takeuchi, 1999), “phosphoserine aminotransferase 1”(Saito et al., 1997), and most frequently “prostate specific antigen” (Woolf-King et al., 2016). There are also many other uses of “PSA” outside of the context of human genes such as “Psoriatic Arthritis” (Schoels et al., 2015) or “Pressure Sensitive Adhesive” (Czech et al., 2013).

The challenge of structuring the knowledge represented in free text is often referred to as “information extraction”, which in turn can be divided into three subtasks (Mooney and Bunescu, 2005). First, “named entity recognition” (NER) is the process of identifying the key concepts that are mentioned in the text. For example, named entities in biomedical texts might include genes, proteins, diseases, and drugs. Second, “normalization” is the application of standardized vocabularies to deal with synonymous concept terms. For example, mentions of “MNAR”, “PELP1, “proline, glutamate, and leucine rich protein 1” would be mapped to the same gene or protein rather than be treated as different entities. Lastly, “relationship extraction”, in which the relations between entities are characterized.

Currently, the gold standard for information extraction in biomedical research is manual review by professional scientists, a process referred to as biocuration (Krallinger et al. 2015). Although there is an active natural language processing community devoted to computational methods for information extraction (Campos, Matos & Oliviera 2012) (Torii et al. 2015) (Usie et al. 2015), the output of these methods are generally not of sufficient quality to be widely used without subsequent expert review.

Previously, we demonstrated that crowdsourcing among non-experts could be an effective tool for NER. We used Amazon Mechanical Turk (AMT) to recruit and pay non-scientists to identify disease concepts in biomedical article abstracts (Good et al. 2015). As a gold standard, we compared AMT workers to professional biocurators who performed the same task (Doğan, Leaman & Lu 2014). We found that, following statistical aggregation, crowd annotations were of very high accuracy (F-score = 0.872, precision = 0.862, recall = 0.883), comparable to professional biocurators.

Although this previous study demonstrated that non-scientists are capable of performing biocuration tasks at a high level, exhaustively curating the biomedical literature using a paid system like AMT is still cost-prohibitive. Citizen science has been successfully applied to the field of biomedical research, but its application in this field has primarily focused on image processing (e.g. Eyewire, Cell Slider, Microscopy Masters), sequence alignment (e.g. Phylo), or molecular folding (e.g. Foldit, EtRNA). Citizen science has also been applied to address language problems, but these are generally focused on transcription (e.g. Smithsonian Transcription Center projects, Notes from Nature, Ancient Lives, reCAPTCHA), translation (e.g. Duolingo), or cognition (e.g. Ignore That, Investigating Word Modalities, Verb Corner). Here, we explored the use of volunteer-based citizen science as a scalable method to perform NER in the biomedical literature. We developed a web-based application called Mark2Cure (http://mark2cure.org) to recruit volunteers and guide them through the same biomedical NER task as we explored in our prior AMT work (Good et al. 2015).

A. In this paper we fulfill our objective of demonstrating that citizen science can successfully be used to address big data issues in biomedical literature. Specifically, we (1) provide a brief overview on the platform we built to enable citizen scientists to do disease NER; (2) demonstrate that citizen scientists are willing to perform the task and inspect our target audience by analyzing the recruitment, retention, and demographics of our participants; (3) demonstrate that citizen scientists are able to perform biocuration tasks when properly trained by assessing the performance of citizen scientists in the same disease NER task used in our AMT experiment; and (4) evaluate the success of our platform using the ‘elements of citizen science success matrix’ developed by Cox et al., and provide our results as a comparison point for other citizen science efforts.

## III. Methods

### A. Document selection

In total, this experiment describes the annotation of 588 documents. These documents were drawn from the training set of the NCBI Disease corpus, a collection of expert-annotated research abstracts for disease mentions (Doğan, Leaman & Lu 2014). Gold standard annotations for 10% of the document set (gold standard documents) were used to provide feedback and were randomly interspersed.

### B. Mark2Cure Design

Mark2Cure was designed to provide a user-friendly interface for engaging members of the public to perform the biocuration task of NER. The goal of this experiment was to have all 588 documents annotated for the NER task by at least 15 volunteers. This threshold was chosen to allow for direct comparison with the results of the AMT experiments. This study and the subsequent survey was reviewed by and approved by the Scripps Health Institutional Review Board and placed in the ‘exempt’ risk category.

At the time of the study detailed in this paper, Mark2Cure was composed of (1) a training module, (2) a feedback interface, (3) a practice module, and (4) a central ‘dashboard’ that organizes volunteer work into a series of ‘quests’. Mark2Cure has always been an open source project: https://github.com/SuLab/mark2cure, and is now being developed to explore additional biocuration tasks.

#### Training

In Mark2Cure, training was a series of four short, interactive tutorials. Training 0 introduced the basic web interface for highlighting concepts in text. Training Steps 1 – 3 introduced the annotation rules distilled from the NCBI disease corpus annotation instructions (Doğan, Leaman & Lu 2014), gradually increased the complexity of the text, and introduced a feedback mechanism to inform the user of their performance (figure S1). The tutorials were designed to provide enough guidance for the participant to perform the task well, with the constraint that overly lengthy tutorials would likely discourage participants. In total there were four tutorials.

#### Feedback

After a user submitted their annotations, feedback was provided by pairing the user with a partner. For visual comparison, Mark2Cure showed the user’s own markings as highlights and their partner’s markings as underlines (figure S2). A score was also calculated and shown based on the F-score (see data and analysis) multiplied by 1000. If the document was designated a gold standard document, then the user’s partner was the gold standard annotations personified as a single ‘expert user’. The gold standard annotations generated by Doğan et al. were attributed to a single ‘expert user’ to facilitate learning by providing gold standard annotations in a recognizable manner consistent with our feedback mechanism. If the document was not a gold standard and no other user had previously annotated the document, then the user was not shown any feedback and was give the full allotment of 1000 points. In all other cases, the user was randomly paired with a user who had previously annotated the document.

#### Practice

Once users completed the tutorials, they were required to work on the practice quest consisting of four abstracts in order to unlock the remaining quests. Users completing the practice documents were always paired with the ‘expert user’ for each of these documents.

#### Quests

For the purpose of organizing documents into manageable units of work, the full set of 588 documents was binned into 118 quests of up to five abstracts each. In addition to the per-document point scoring system (described previously), a quest completion bonus of 5000 points was awarded upon completion of all five abstracts. For the sake of usability, users who started a quest were allowed to finish it even if the quest was subsequently completed by the community. Hence it is possible for abstracts to be completed by over 15 different users.

### C. Data and Analysis

Mark2Cure was set up with Google Analytics for site traffic analysis. During the experiment period, emails were sent weekly to the participants via Mailchimp, and Mailchimp analytics provided open and click rate information. Mark2Cure also logged information regarding user sign ups, training and submissions. Precision, recall, F-scores, data quality and cost metrics were calculated as previously described (Good et al. 2015). A survey was sent via email to the subscribers on the Mark2Cure mailing list at the end of the experiment (379 subscribers), and 78 of the subscribers (20.5%) responded to the survey. The metrics developed by Zooniverse were loosely applied to study the success of the project, though these metrics are better suited for project suites with multiple longer-running projects. The active period for this experiment started on January 19^th^ and ended on February 16^th^ 2015. An export of the data generated in this experiment (which was used for the analysis) can be found at: https://figshare.com/articles/Mark2Curator_annotation_submissions_for_NCBI_disease_corpus/2062554

## IV. Results

### A. Training and Retention

Training 0 introduced the highlight mechanism underlying Mark2Cure. Training 1 introduced rules for highlighting disease mentions in a sentence-by-sentence manner. Training 2 allowed users to practice the rules they had learned while acclimating the user to longer spans of texts. Training 3 introduced the feedback and scoring mechanisms. User tests prior to the launch of this experiment indicated that all training steps could be completed in under 20 minutes, but user drop off had not been determined at that stage.

Based on Mark2Cure’s log files, 331 unique users completed training 1, 254 unique users completed training 2, and 234 unique users completed training 3. User drop off was highest between Training 1 and Training 2, but 92% of users that completed Training 2 went on to complete Training 3. Of the 234 unique users that completed the training, over 90% of the users (212) contributed annotations for at least one document. To better understand the user drop off and retention throughout the different training pages, we obtained the unique page views and average page view time for each training page using Google Analytics (figure S3). Problem pages within the tutorials identified with Google Analytics were confirmed by emails received from users having trouble on those pages.

Beyond the training modules, Mark2Cure participants were paired with other users and given points based on their performance in order to encourage continuous learning/improvement. While this was effective when users were paired with the gold standard ‘expert user’, many users expressed frustration when paired with poor performing partners. Issues with partner pairing highlighted the need to apply sorting mechanisms or allow for ‘expert trailblazers’ like those utilized in Eyewire.

In addition to responding to each email received, Mark2Cure published 204 tweets, nine blog posts, and sent eight newsletters during the 28 days that this experiment was running. We estimate having 466 email interactions with the users and 221 other communications during this time yielding values of 0.282 and 0.594 for our communications and interactions metrics respectively (table 1B). Post-survey results indicated that user interface issues were a common (but generally surmountable) obstacle to participation.

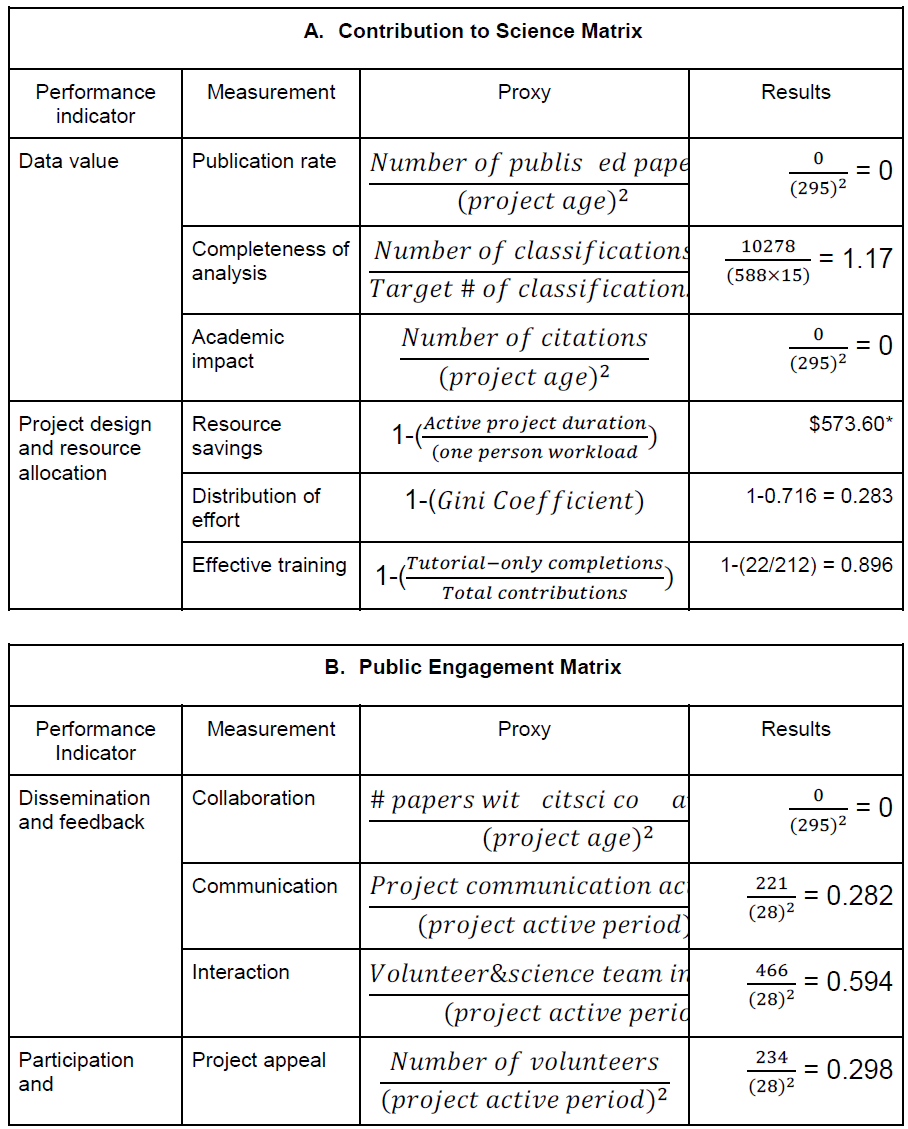

**Table I.**
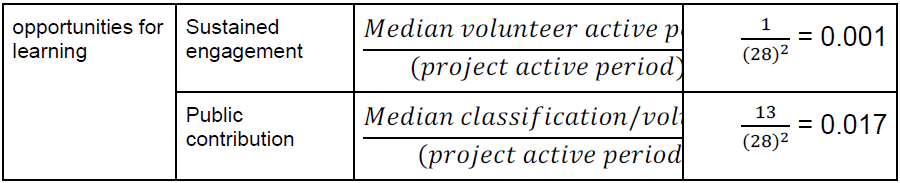
Computed metrics of “Contributions to Science” and “Public Engagement”, as defined by the Zooniverse projects (Cox et al. 2015). As a single citizen science project, we cannot perform the internal comparisons used in the Zooniverse paper and thus have no basis for comparing our results (Zooniverse results are normalized). Hence, we provide non-normalized results which may serve as a basis for comparison for other research teams who also only have one citizen science project. The resource savings value was not calculated based on traditional workload, as we already had Amazon Mechanical Turk experiments demonstrating the cost to complete the experiment (though the original cost of generating the NCBI Disease Corpus via professional biocuration is not known). In terms of Sustained Engagement, the “median volunteer active period” was based on dates of account creation and time stamp of their last submission. Because many contributors participated just once, the median active period was one day for this experiment, resulting in a sustained engagement of 0.001.

### B. Recruitment, Analytics, and Demographics

Five months prior to the launch of this experiment, we began to blog periodically about Mark2Cure and engage with the rare disease community on Twitter. We focused on the rare disease community because many members of this community are highly motivated to read scientific literature, engaged in research, active on social media channels, and experienced with outreach. By the time this experiment was launched, we had a mailing list of 100 interested potential users and 75 followers on Twitter; however, only about 40 users signed up after the first 10 days of the experiment’s launch.

Sessions from new users peaked the day after the article on Mark2Cure was published in the San Diego Union Tribune (figure 1). A second, smaller surge in sessions from new users was observed the day after a blurb on Mark2Cure was published on California Healthline; however, total sessions (from both new and returning users) rivaled that of the one seen from the San Diego Union Tribune article.

**Figure 1:**
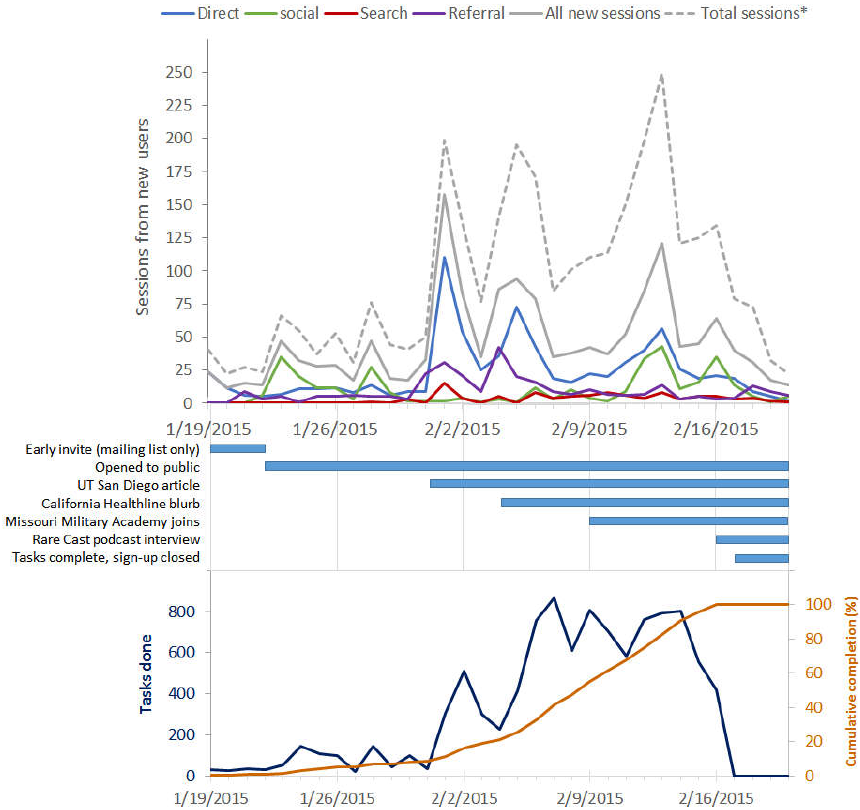
Top: Google Analytics of Mark2Cure’s new-user sessions broken down by the source of the sessions. *Total sessions include sessions from both new and returning users. Middle: Timeline of significant events throughout this experiment. Bottom: The number of tasks done on a daily basis (dark blue) along with the cumulative tasks done as a percent of total completion (orange).

New users from social media peaked around Feb. 13^th^ and 14^th^, as existing users were actively marking documents, tweeting about their activities, encouraging other users to do the same, and luring new users to try Mark2Cure. Over the course of this experiment, one user posted over 50 tweets and generated Mark2Cure specific hashtags to encourage others to join the effort. New users from social media peaked again on Feb. 16^th^ with the release of Global Gene’s RareCast podcast interview featuring Mark2Cure. As with many large rare disease communities and organizations, Global Genes has a strong social media presence.

Survey results at the end of the experiment indicated that 67% of respondents learned about Mark2Cure from the newspaper, 24% learned about Mark2Cure from social media channels, while 15% learned about Mark2Cure from a friend or a google search (figure 2C). In addition to recruiting via traditional press and social media channels, recruitment was also driven by members of the NGLY1-deficiency community. Participants from this rare disease community made a concerted effort to ensure that this experiment would be completed successfully, with the hopes that Mark2Cure could be applied to accelerate research on NGLY1-deficiency. One participant from this community approached the Missouri Military Academy (MMA) and recruited both instructors and students to participate. Over 24 participants from MMA contributed about 10% of the task completions.

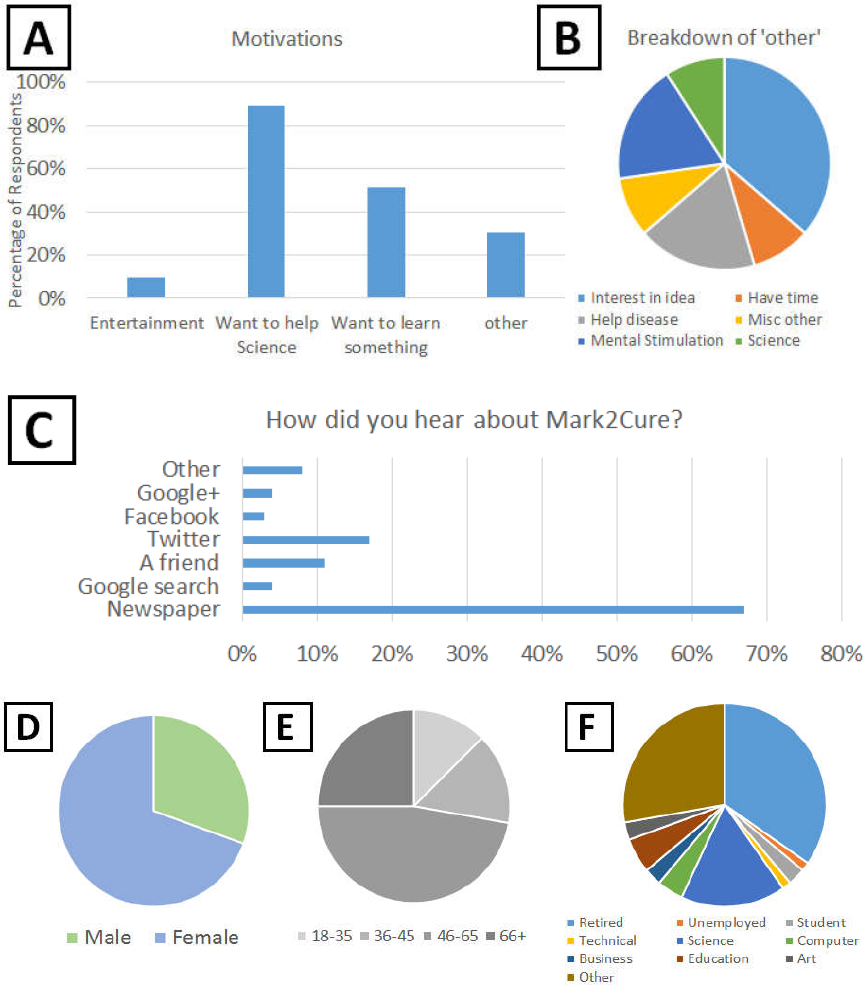

**Figure 2.**
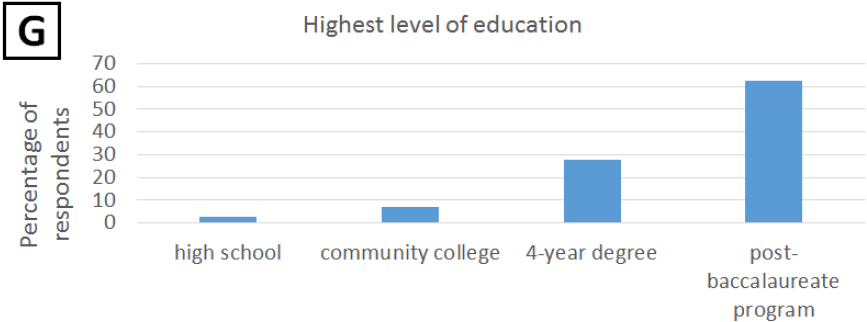
– User survey results. A. Non-exclusive motivations for participating. Users could select from a list of categories used in the AMT experiments OR enter a free-text response. 89% of the end survey respondents wanted to help science, 51% wanted to learn something, and 10% were looking for entertainment. B. Further analysis of the ‘other’ motivations for participating. C. How participants discovered Mark2Cure. D. Ratio of female to male survey respondents was 69% to 31% respectively. E. Age demographics of the survey respondents. 28% of respondents were 18-45 years old, and 72% were 46 years of age or older. F. Occupational fields of the survey respondents. In terms of occupations, 35% of respondents were retired, 22% worked in a science, computer or technical field; 28% were care providers, science communicators, or journalists. Only 4% of respondents were students or unemployed, and the remaining 11% of respondents were employed in business, education, or art. G. Educational distribution of survey respondents. 83% of contributors completed a four-year college degree or higher.

The majority (65%) of survey respondents cited the ‘desire to help science’ as their motivation for participating (figure 2A). Though not necessarily representative, the results of our survey suggest that the participants in this experiment were demographically quite different from our AMT experiments. Women were more likely to participate (or report their participation) in our survey than men (figure 2D). On average, our participants were older than the participants from the AMT experiments (figure 2E), in part due to the readership/recruitment from the San Diego Union Tribune article (figure 2C). After the publication of that article, we received many inquiries about participating in the project from citizen scientists who volunteered information about their employment status and age (particularly from retirees). In pooling the contributions from just 14 of the 212 participants (6.6%) who volunteered this information (without ever being asked), we found that seniors and/or retirees contributed at least 26% of total document annotations in this experiment. In our demographic survey, 18 respondents (25%) reported being 66 years of age or greater (figure 2E), and 25 respondents (35%) reported being retired (figure 2F); hence, the actual contributions from this demographic group are likely to be higher. Student participation was underreported (3%) in our survey and does not reflect the concerted effort of students from the Missouri Military Academy.

### C. Distribution and Performance

Over the 28-day experiment, 212 users submitted 10278 annotated abstracts with over half of the annotated abstracts submitted in the final week of this experiment. As with all crowdsourcing systems, the distribution of task completions among users was skewed. There were 22 users who completed over 100 documents, 92 who completed over 10, and 98 who completed 10 or less (figure 3A). Roughly 80% of the document completions were submitted by 24% of the contributors, more similarly following the pareto principle than the 90-9-1 rule as Mark2Cure was not setup for interaction/discussion between users at this point in time.

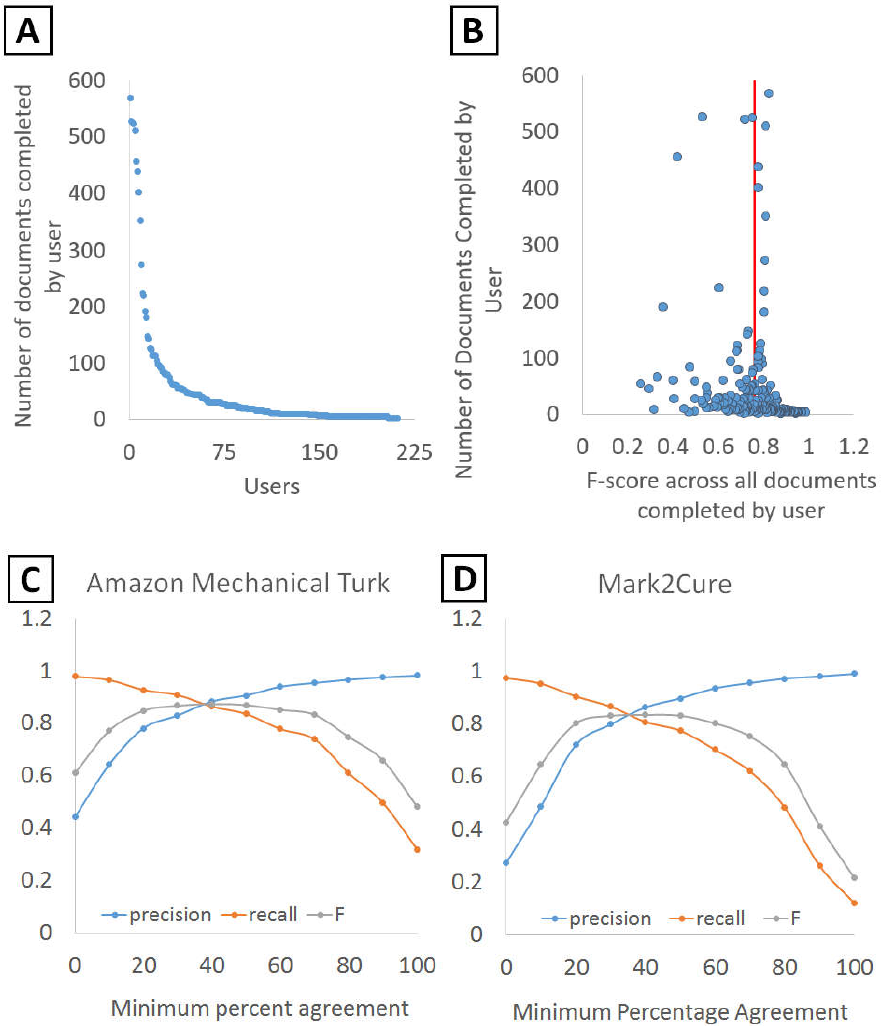

**Figure 3.**
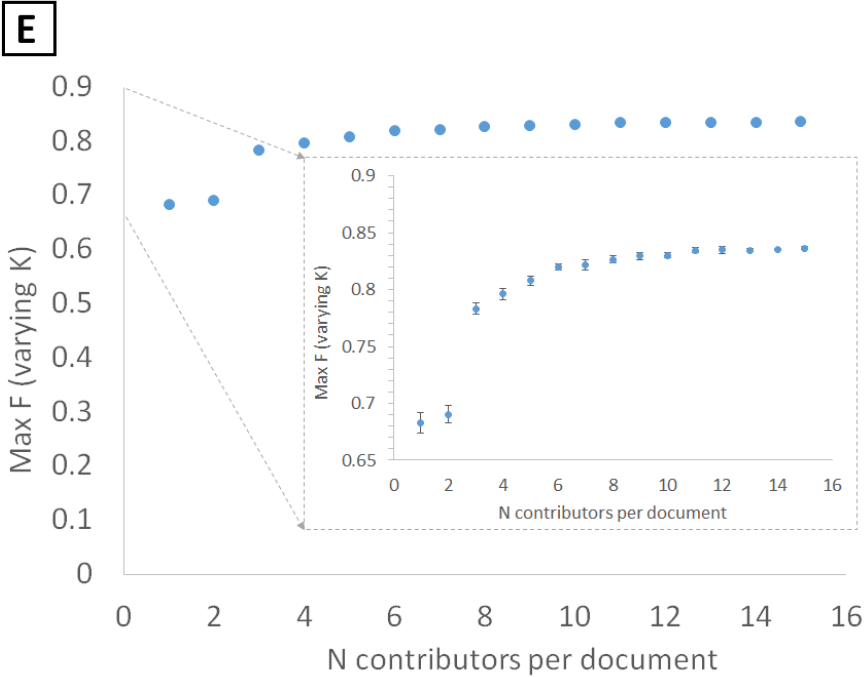
– A. The number of documents processed per user. B. Quality of each user’s contributions based on the number of documents that user completed. Each user’s F-score was calculated based on their contributions across all of the documents. The average user F-score is indicated by the red line. C. Effects of the minimum percentage agreement between annotators on the level of agreement with the gold standard in the Amazon Mechanical Turk experiments and in this experiment for Mark2Cure (D). E. The impact of increasing the number of contributors per abstract on the quality of the annotations.

Overall, the accuracy of contributions relative to the gold standard was quite high. The average F-score per user across all of their annotations was 0.761 with a standard deviation of 0.143 (figure 3B), which was on par with that of our previous AMT results (Good et al. 2015).

To assess the aggregate accuracy across all users, we computed the “minimum percent agreement” for each annotation, defined as the number of users marking a given annotation divided by the total number of users to process the document (usually 15), and computed accuracy statistics at multiple thresholds (Figure 3D). At 0% minimum agreement (taking the union of all user annotations), we observed precision of 0.274 and recall 0.976. At 100% minimum agreement (the intersection of all user annotations), we observed precision of 0.992 and recall of 0.122. The maximum F-score of 0.836 was reached at 40% minimum agreement.

These accuracy results were also very similar to our AMT experiments (figure 3C), in which precision ranged from 0.444 to 0.983 and recall varied from 0.980 to 0.321 (at a 0% and 100% agreement threshold, respectively). The maximum F-score of 0.875 was also reached at a minimum agreement threshold of 40%. For comparison, these aggregate maximum F-scores were also on par with the individual expert annotators who performed the initial phase of annotations for the disease corpus (Doğan & Lu 2012).

The difference in maximal F-score could be attributed to the differences in the way the rules were presented in AMT vs Mark2Cure, or due to the difference in incentive structures and goals. In AMT, poor performers can be blocked or may self-select out of a task so as to not affect their ability to qualify for future tasks (poor performance reports can affect an AMT workers ability to qualify for jobs). In contrast Mark2Cure contributors are never blocked and are encouraged to continue contributing as data quality is only one of several metrics by which success is determined. Overall, the performance by citizen scientists was on par with that of AMT workers on this task, even though the demographics of the citizen scientists were quite different from those of our AMT workers.

To assess how increasing the number contributors affected the quality of the aggregate annotations, we simulated smaller numbers of annotators per document by randomly sampling from the full dataset. We found that the greatest increase in F-score was observed when the number of annotators was increased from two (F=0.690) to three (F=0.783) (figure 3E).

Qualitative inspection of user disagreement in the annotations revealed issues with conjunction highlighting for certain users. For example, terms like ‘colorectal cancer’ would be highlighted individually as ‘colorectal’ and ‘cancer’. It is unclear whether or not this type of error reflected the user’s understanding of the rules or if the user had issues with the highlighting function, as there were reported difficulties in highlighting conjunctions spanning more than one line. Another major source of disagreement was the inclusion of modifying terms. Eg– The inclusion of ‘premenopausal’ in ‘premenopausal ovarian cancer’. This type of disagreement was unsurprising given the inconsistent inclusion of these modifiers in the gold standard. For example, ‘early-onset’ is a modifying term that is inconsistently included with ‘breast cancer’ in the gold standard. Some gold standard documents will have the include the entire term ‘early-onset breast cancer’, whilst others will only have ‘breast cancer’. This reflects an area in the annotation rules that could be improved by biocurators in the original data set.

### D. Elements of Successful Citizen Science Matrix

Data quality has been the focus of much academic research on citizen science, but it is not the only measure of success. Many citizen science projects have additional goals of engaging people in science and motivating them to incorporate scientific thought, hence the process of engaging citizen scientists can in itself also be a measure of success (Freitag & Pfeffer 2012).

As one of the most established metrics, data quality is the most uniformly examined. Recent efforts have been made to define, expand, and apply additional metrics of citizen science success. The Zooniverse project created a set of useful metrics that could be applied across many citizen science projects. These metrics were normalized internally so that different projects by the Zooniverse team can be compared with one another. Although these metrics can be very useful for internally evaluating different projects, the normalization used in the Zooniverse paper means that that the reported results cannot be used for comparison purposed by researchers with only one citizen science project. Hence, we calculated and reported all the Zooniverse metrics as one data point to which other citizen science projects can be compared (Table 1).

As this is the first publication about Mark2Cure, the performance metrics based on citations and publications such as Publication Rate, Academic Impact, and Collaboration are not meaningful. For the Completeness of Analysis metric, Mark2Cure performed well; however, this reflected more on the flexibility of the project period--remaining open until all the data was collected. Hence, this metric might not be as useful for projects with flexible timelines that open and close based on the data needed and the data collected. In spite of the flexible timeline, this phase of Mark2Cure was rather short and ended just as recruitment improved, which may explain the low Sustained engagement result. Mark2Cure scored well in Interaction and Effective training which actually reflects difficulties users had with Mark2Cure. Much of the interaction initiated by users was due to problems with the tutorials or interface; and by addressing these issues quickly we were able to encourage many users to complete the tutorial and make a meaningful contribution. The Distribution of effort was higher than Zooniverse’s across project average of 0.18 (Simmons 2015); but similar to the Andromeda Project, this higher Distribution of effort score may be an artifact of the short project period.

## V. Discussion

In the 1980’s, an information scientist named Don Swanson found that several abstracts about dietary fish oil contained mentions of blood viscosity, platelet function, and vascular reactivity. Swanson also found the same terms in abstracts from a disparate body of literature surrounding Raynaud’s syndrome, allowing him to uncover the relationship between Raynaud’s syndrome and fish oil—an undiscovered relationship even though all the information to establish the link was already publicly available (Swanson 1986). This hidden knowledge was uncovered by Swanson at a time when the biomedical literature was growing at an annual rate of about 10,000-15,000 articles. The rate of biomedical literature publication now exceeds 1 million articles per year and represents a body of knowledge that is increasingly difficult to harness. Information extraction is a necessary step towards harnessing the undiscovered but already available knowledge; however, it encapsulates some of the most time-consuming tasks in biocuration.

As the number of professional biocurators shrinks relative to the volume of literature to be curated, alternative strategies for keeping pace need to be explored (Howe et al. 2008). The entrance of citizen science into this domain opens up many new opportunities. Most immediately, citizen scientists can help to generate new annotated corpora for training and evaluating computational methods for information extraction (Good et al. 2015). Following Galaxy Zoo’s example (Richards & Lintott 2012), we can set up computational systems that learn to perform the current tasks of the citizen scientists. Once these methods reach acceptable levels of performance, the citizens can be directed towards other areas still in need of human input.

We were fortunate that media attention allowed us to recruit enough participants to complete this phase of the project; however, recruitment and sustained engagement (retention) remain an important issue for Mark2Cure. Many citizen science projects (Zooniverse, Eyewire, Foldit) have demonstrated that recruitment improves as results are produced, and it makes no sense for our small citizen science community to attempt to tackle volumes of literature too large to complete or see results. In order to grow at a sustainable pace, Mark2Cure is currently focused on NER of three concept types in abstracts surrounding NGLY1 deficiency--the rare disease that was of interest to the greatest number of our participants during the early phase of Mark2Cure. By focusing on literature in a specific disease domain (especially one of interest to previous participants), we reward organized participation from patient/community groups to encourage recruitment and narrow the range of literature to a volume that is manageable by the citizen science community. Although we currently apply Mark2Cure to create an annotated, NGLY1- deficiency-specific corpora, we are also developing Mark2Cure towards more challenging areas of information extraction such as relationship extraction. We expect extracting relationship information in NGLY1-related literature to provide more insightful information and be of greater utility to the NGLY1 researchers, ultimately leading to new discoveries in this field.

Apart from helping to develop computational methods for information extraction, citizen scientists can help at much higher levels than machines are likely to reach. A motivated community of citizen scientists can accomplish nearly any goal, including the development of their own computational methods for solving complex tasks (Khatib 2011). Within the domain of biocuration, citizen scientists could help with challenges like: prioritizing the most interesting documents for curation, developing controlled vocabularies, annotating images in documents, creating summaries, or even soliciting funding for research. The scientific possibilities are limitless. To achieve them will require the development of strong synergistic cooperative relationships. In addition to large numbers of participants, experts in information extraction who can help dramatically increase the efficiency of processes for integrating that knowledge need to cooperate with professional biomedical scientists who can turn the scattered information in the public domain into new knowledge. One caveat common to many domains of research where citizen science has been applied is the reluctance of researchers to collaborate due to of ‘data quality concerns.’ We accounted for researcher cooperation when selecting the scope of our project’s current efforts, and hope to demonstrate with current and future iterations of Mark2Cure how large communities of citizens whose health and well-being will ultimately benefit from their work can become fundamentally important contributors to the production of a ‘problem solving ecosystem’ (Michelucci 2016).

## VI. Conclusion

As a citizen science project, Mark2Cure would be classified as a science-oriented virtual project (Wiggins & Crowston 2011) subject to the issue of ensuring valid scientific results while designing for online participation/interest. We addressed data quality issues as other citizen science opportunities have, by using replication across multiple participants, having participants evaluate established control items, and using a corpus of text that had already been expertly reviewed as a benchmark (Wiggins & Crowston 2014). As Wiggins (2014) pointed out, data quality issues in citizen science are often project design issues; hence data quality can often be improved by analyzing participant interaction and adjusting the design. By using the gold– standard NCBI disease corpus (as we did in the AMT experiments) and formulating our tutorials around the annotation rules set forth in the development of that corpus, we demonstrate that citizen scientists are willing to perform NER tasks and (in aggregate) can perform comparably with expert curators. Furthermore, we demonstrate how researchers might utilize site traffic information and logged data in order to improve aspects of the design in order to achieve quality data, and we analyze our project with accordance to Cox et al’s ‘Elements of citizen science success matrix’ providing a comparison point for other citizen science efforts.

## VI. Acknowledgements

This paper would not be possible without the citizen scientists who participated in this experiment, and continue to contribute to our current efforts at http://mark2cure.org. We thank the members of the NGLY1-deficiency disease community who helped with our recruitment efforts and the faculty, staff, and student participants from the Missouri Military Academy. We especially thank Bradley Fikes of the San Diego Union Tribune for his excellent piece on Mark2Cure, which helped immensely in our recruiting efforts.

Contributors who gave us permission to publish their names can be found here: http://mark2cure.org/authors/beta_experiment

This work was supported by the US National Institute of Health (U54GM114833 to A.I.S.). This work was also supported by the Scripps Translational Science Institute, an NIH-NCATS Clinical and Translational Science Award (CTSA; 5 UL1 RR025774).

